# A high-sensitivity ELISA for detection of human FGF18 in culture supernatants from tumor cell lines

**DOI:** 10.1101/2023.06.16.545249

**Authors:** Yuichi Tsuchiya, Sachiko Komazawa-Sakon, Masato Tanaka, Takumi Kanakogi, Kenta Moriwaki, Hisaya Akiba, Hideo Yagita, Ko Okumura, Kevin C Entzminger, CJ Okumura, Toshiaki Maruyama, Hiroyasu Nakano

## Abstract

Fibroblast growth factor 18 (FGF18) is elevated in several human cancers, such as gastrointestinal and ovarian cancers, and stimulates the proliferation of tumor cells. This suggests that FGF18 may be a promising candidate biomarker in cancer patients. However, the lack of a high-sensitivity enzyme-linked immunosorbent assay (ELISA) does not permit testing of this possibility. In this study, we generated monoclonal antibodies against human FGF18 and developed a high-sensitivity ELISA to measure human FGF18 at concentrations as low as 10 pg/mL. Of the eight tumor cell lines investigated, we detected human FGF18 in culture supernatants from four tumor cell lines, including HeLa, OVCAR-3, BxPC-3, and SW620 cells, albeit the production levels were relatively low in the latter two cell lines. Moreover, the in-house ELISA could detect murine FGF18 in sera from mice overexpressing murine *Fgf18* in hepatocytes, although the sensitivity in detecting murine FGF18 was relatively low. This FGF18 ELISA could be a valuable tool to validate FGF18 as a potential biomarker for cancer patients and to test the contribution of FGF18 for various disease models in vivo and in vitro.

## 1. Introduction

Fibroblast growth factors (FGFs) constitute 22 family members and are involved in various cellular responses, including organogenesis, wound healing, fibrosis, proliferation, and carcinogenesis [1–3]. FGFs are assigned to three groups depending on their mode of action as follows: intracrine, paracrine, and endocrine factors [2,3]. Intracellular FGFs are not secreted and do not interact with FGF receptors (FGFRs), thereby acting in the cells. Paracrine FGFs, including 15 members of the FGF family, are further divided into seven subfamilies and act locally due to their high affinity for heparan sulfate proteoglycans that prevent diffusion through the extracellular matrix. Thus, paracrine FGFs work locally near the place of secretion. In contrast, endocrine FGFs, including FGF15/19, FGF21, and FGF23, are released from cells and circulate in the blood. Hence, endocrine FGFs, like hormones, exhibit their action on distant sites. Most FGFs bind to FGFRs 1 to 4, which contain tyrosine kinase activity and initiate the PKC, AKT, and MAPK pathways [4,5].

FGF18 belongs to the FGF8 subfamily of paracrine FGFs, including FGF8, FGF17, and FGF18, and plays a crucial role in the development of osteogenesis, chondrogenesis, and lung alveolar development [6–8]. Regarding tumor development, many studies have reported that the expression of FGF18 is elevated in different cancers, including colorectal, ovarian, and gastric cancers [9]. Sonvilla et al. showed that FGF18 expression is elevated in colorectal tumors and promotes tumor cell growth by enhancing the proliferation of colon-associated fibroblasts as well as tumor cells and neovascularization [10]. Wei et al. reported that FGF18 is overexpressed in advanced-stage, high-grade serous ovarian cancer, and its elevation is correlated with poor survival [11]. These results suggest that FGF18 is a prognostic and therapeutic biomarker in some cancer patients. However, as far as we have investigated, there is no reliable and highly sensitive enzyme-linked immunosorbent assay (ELISA) to detect human FGF18. Thus, the development of an FGF18 ELISA is crucial for testing FGF18 as a potential biomarker to diagnose or predict the progression of tumors in some cancer patients.

We developed a high-sensitivity ELISA for human FGF18 (hFGF18) by generating monoclonal antibodies against hFGF18. Our in-house ELISA detected hFGF18 at concentrations as low as 10 pg/mL in culture supernatants from several cancer cell lines, such as HeLa, OVCAR-3, and BxPC-3. Moreover, we detected murine FGF18 (mFGF18) in sera from mice overexpressing murine *Fgf18* in hepatocytes, although the sensitivity in detecting mFGF18 by ELISA was relatively low. Together, the FGF18 ELISA system could be a valuable tool to validate FGF18 as a potential biomarker for cancer patients and test the contribution of FGF18 to various disease models in vivo and in vitro.

## 2. Materials and Methods

### 2.1. Reagents and cell culture

Human FGF18ΔC8 (Peprotech 100-28) and full-length murine FGF18 (ProSpec CYT-064) were purchased from the indicated sources. Anti-hFGF18 antibody (CSB-PA120525) was purchased from Cusabio. Streptavidin poly-HRP80 conjugate (65R-S119) was purchased from Stereospecific Detection Technologies GmBH.

HeLa, OVCAR-3, BxPC-3, PANC-1, Huh7, SW620, and HT29 cells were obtained from ATCC. FLC4 cells were provided by E. Miyoshi. All cell lines were maintained in DMEM containing 10% fetal bovine serum.

### 2.2. Mice

The generation of *Fgf18* Tg mice that express murine *Fgf18* in hepatocytes will be published elsewhere. All experiments were performed according to the guidelines approved by the Institutional Animal Experiments Committee of Toho University School of Medicine (19-409 and 19-412).

### 2.3. Generation of a mammalian expression vector for hFGF18 and purification of hFGF18

hFGF18 cDNA was amplified by RT‒PCR using mRNAs from HepG2 cells as a template. The amplified cDNA of hFGF18 was subcloned into the C-terminal FLAG Tag vector, pcDNA3-FLAG (provided by T. Natsume), and resulted in the generation of pcDNA3-hFGF18-FLAG. The following primers were used to amplify the cDNA of hFGF18: forward, 5’-GCGCG<u>AAGCTT</u>GCGATGTATTCAGCGCCCTCCGC-3’ and reverse, 5’-GCGCG<u>AGATCT</u>GGCAGGGTGTGTGGGCCGGATCC-3’. HEK293 cells (4 × 10^6^ cells/150 mm dish) were transiently transfected with pcDNA3-hFGF18-FLAG using PEI-MAX (Polysciences 24765). The culture medium was changed 24 hours after transfection, and transfected cells were cultured for an additional 3 days. The culture supernatants were collected, pooled from 50 x 150 mm dishes, and used for subsequent steps.

We directly applied the culture supernatants to heparin columns (GE Healthcare GE17-099-801) to purify hFGF18 secreted in the culture supernatants from HEK293 cells. Then, the column was extensively washed with 10 mM Tris-HCl pH 8.0 containing 0.5 M NaCl, and hFGF18 was eluted with 10 mM Tris-HCl pH 8.0 containing 1.0 M NaCl. The purified hFGF18 was dialyzed in PBS and used for immunization.

### 2.4. Generation of antibodies against human FGF18

To generate rat anti-hFGF18 antibodies, we subcutaneously immunized Wistar rats in the footpads 4 times with purified hFGF18Δ1C emulsified in TiterMax Gold (TiterMax). Three days after the final immunization, bilateral inguinal and popliteal lymph node cells were fused with NSObcl2 myeloma cells using PEG1500 (Roche) [12]. Hybridoma cells were selected in DMEM/10% FCS containing HAT (Sigma, MO) and 1% BM-Condimed (Roche). One of the hybridomas (clone 12G7-9) that showed high reactivity was used to generate an ELISA. Large-scale antibody production was performed using Hybridoma SFM (Gibco 12045076) and CELLine 1000 bioreactors (WHEATON). Antibodies were purified from the culture supernatant using caprylic acid and dialyzed against PBS.

To generate rabbit anti-hFGF18 antibodies, we immunized rabbits with purified hFGF18Δ1C four times at 2-week intervals using complete Freund’s adjuvant for the first injection and incomplete Freund’s adjuvant thereafter. The messenger RNA was purified from the total RNA of the spleen and bone marrow cells of immunized rabbits using the NucleoTrap mRNA kit (Macherey-Nagel, 740655) and according to the manufacturer’s protocol. Then, variable regions of heavy and kappa chains of immunoglobulin genes were amplified by PCR (US patent number 9,890,414) and subcloned into the phage display vector, resulting in phage library generation. After four rounds of panning to enrich phage clones that specifically bound to precoated hFGF18ΔC, isolated phages were used to infect *E. coli* cells and to prepare single colonies for screening individual clones. Ninety-five single colonies were picked from each library and cultured to produce soluble Fab fragments in the presence of 1 mM isopropyl β-d-1-thiogalactopyranoside (IPTG). Then, the reactivity of each Fab fragment to hFGF18 was determined by ELISA. Both heavy and kappa chains from clones showing strong binding activities were sequenced. The variable regions of heavy and kappa chains were inserted into a modified pTT5 bicistronic IgG expression vector (National Research Council, Canada). HEK293 cells were transfected with the bicistronic vector encoding heavy and kappa chains of rabbit IgGs, and IgGs were then purified from the culture supernatants by Protein A-column chromatography (GE Healthcare, 17-1279-01). We obtained 6 clones as follows: 19B-SH1, 19S-SD1, 19S-SE5, 19S-SH1, 19S-SH4, and 19S-SH7.

The biotinylation of these antibodies was conducted using the EZ-Link Sulfo-NHS-LC-Biotin No-Weigh Format (Thermo A39257).

### 2.5. Generation of C-terminal deletion mutants of hFGF18 and mFGF18

A C-terminal deletion mutant of hFGF18 lacking 25 amino acids, excluding the signal peptides (hFGF18ΔC25), was amplified by PCR using pcDNA3-hFGF18-FLAG as a template. PCR-amplified fragments were subcloned into pGEX-6P-1 (GE Healthcare, 28-9546-48). The following primers were used: forward, 5’-GCGCG<u>CAATTG</u>GTTGCCGAGGAGAACGTGGAC-3’ and reverse, 5’-GCGCG<u>CTCGAG</u>TCACTGCCCCTTGGGGTAGCGC-3’. To generate the full-length and C-terminal deletion mutant of mFGF18 lacking 8 amino acids and excluding the signal peptides (mFGF181ΔC8), RT‒PCR-amplified fragments encoding the fragments were subcloned into pGEX-6P-1. The following primers were used: forward, 5’-CCG<u>CAATTG</u>GCCGAGGAGAATGTGGACTTC-3’ and reverse, 5’-GGCC<u>GCGGCCGC</u>CTAGCCGGGGTGAGTGGGGCG-3’ for full-length mFGF18; forward, 5’-CCG<u>CAATTG</u>GCCGAGGAGAATGTGGACTTC-3’ and reverse, 5’-GGCC<u>GCGGCCGC</u>CTACCGGGATCGCTTGGTGAC-3’ for mFGF181ΔC8 (the underlined sequences are restriction enzyme sites). Mutants were expressed in BL21 as a standard procedure and then purified with glutathione-sepharose (GE Healthcare, 17-0756-01) as previously described [13]. Each GST-fusion protein was released from GST adsorbed to glutathione-sepharose by PreScission protease (GE Healthcare, 27-0843-01). The concentrations of purified proteins were determined by Bradford analysis (Bio-Rad 5000006). The purity of these proteins was evaluated by staining with Coomassie blue.

### 2.6. Detection of plate-coated recombinant FGF18

Purified hFGF18 from HEK293 cells or mFGF18 was serially diluted with PBS, coated onto the plate at 4 °C overnight, and blocked with 10% BSA-PBS-0.05% Tween (PBS-T) for 1 hour. After washing with PBS-T three times, the plate was incubated with biotinylated antibodies as indicated at RT for 1 hour. After washing with PBS-T three times, the plate was incubated and developed with streptavidin-HRP (DAKO P039701-2) and TMB substrate (BD OptEIA, 555214).

### 2.7. Enzyme-linked immunosorbent assay (ELISA)

We generated an in-house ELISA to measure the concentrations of hFGF18 and mFGF18. Briefly, the plate was coated with clone 12G7-9 (30 µg/mL) in 0.5% BSA-PBS at 4 °C overnight and blocked with 1% BSA-PBS-0.05% Tween (PBS-T) for 1 hour. After washing with PBS-T three times, samples were added to the plate and incubated at RT for 1 hour. After washing with PBS-T three times, the plate was incubated with biotinylated 19S-SE5 antibody (1.0 μg/mL) at RT for 1 hour. After washing with PBS-T three times, the plate was incubated with streptavidin poly-HRP80 conjugate (0.2 μg/mL) at RT for 1 hour and developed with TMB substrates. The lower detection limits of hFGF18 and mFGF18 were approximately 10 pg/mL and 500 pg/mL, respectively.

### 2.8. Quantitative polymerase chain reaction assays

Total RNA was extracted from various human tumor cell lines, and cDNA was synthesized with the Revertra Ace qPCR RT Kit (Toyobo). Quantitative polymerase chain reaction (qPCR) analysis was performed with the 7500 Real-Time PCR detection system and the SYBR Green method (SYBR Green PCR master mix, ThermoFisher 4312704) of the target genes together with a human *GAPDH* endogenous control and the use of 7500 SDS software (Applied Biosystems). The primers used in this study were as follows: *FGF18*-F, 5’-TGCTTCCAGGTACAGGTGCT-3’; *FGF18*-R, 5’-GCTGCTTACGGCTCACATCG-3’; *GAPDH*-F, 5’-AGCCACATCGCTCAGACAC-3’; and *GAPDH*-R, 5’-GCCCAATACGACCAAATCC-3’.

### 2.9. Knockdown of FGF18 by siRNAs

HeLa cells were transfected with control or FGF18 siRNAs (Dharmacon SO-2989166G) using Lipofectamine RNAiMAX (Thermo Fisher 137780). The medium was changed 24 hours after transfection, and the cells were cultured for 3 days, and The expression of FGF18 in HeLa cells was analyzed by qPCR. The concentrations of FGF18 in culture supernatants were determined by ELISA.

### 2.10. Measurement of the concentrations of mFGF18 in sera from Non-Tg and Fgf18 Tg mice

Sera were collected from 6- to 8-week-old mice, and the concentrations of mFGF18 were determined by the in-house ELISA.

### 2.11. Statistical analysis

Statistical analysis was performed using the two-tailed unpaired Student’s *t-*test. *P* value < 0.05 was considered significant.

## 3. Results and Discussion

### 3.1 Generation of anti-human FGF18 antibodies

We generated anti-FGF18 monoclonal antibodies to test whether FGF18 may be a biomarker in several cancer patients. We attached the C-terminal FLAG tag to hFGF18 to purify hFGF18 using an anti-FLAG tag antibody from the culture supernatants of HEK293 cells following transient transfection; however, the C-terminal FLAG tag was lost during the translation and secretion of hFGF18. Consistent with this finding, a previous study reported that the C-terminal amino acids of mFGF18 are cleaved by an unknown protease [14]. Moreover, FGF23 is cleaved by one of the proprotein convertases, furin, which recognizes the consensus sequence RXXR↓X (R, arginine; X, any amino acid; ↓, cleavage site) and cleaves the site between the last R and X [15]. Comparing the C-terminal amino acid sequence of hFGF18 to the consensus RXXR motif (Fig. 1a), hFGF18 may be cleaved between the amino acids arginine^200^ and isoleucine^201^ or arginine ^202^ and proline^203^. These results suggest that C-terminal 5 or 7 amino acids may be cleaved during the maturation of hFGF18. However, the precise cleavage site of hFGF18 expressed in mammalian cells was not determined in this study, hence, we tentatively refer to it as hFGF181ΔC purified from HEK293 cells.

**Fig. 1.**
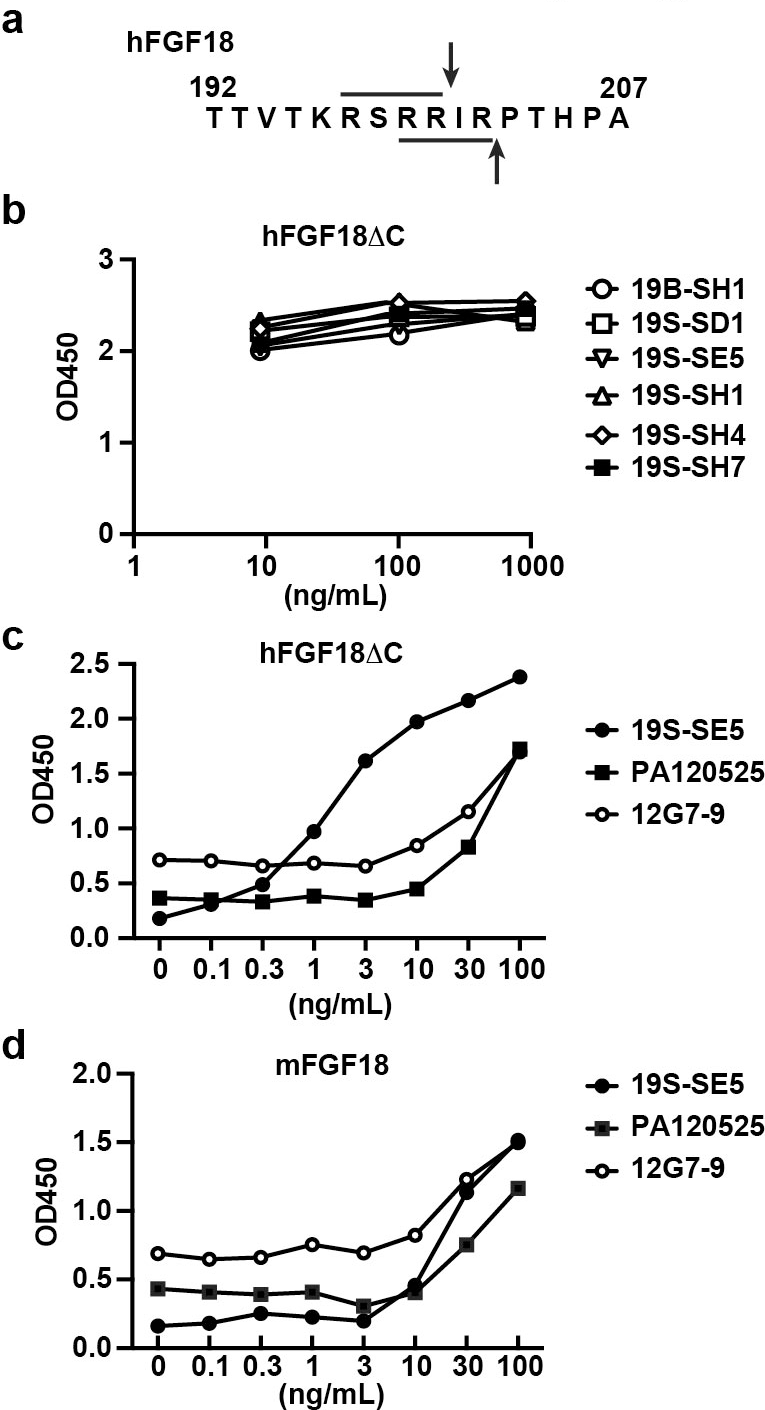
Generation of anti-human FGF18 antibodies. **a,** Alignment of the consensus motif recognized by furin (RXXR) and the C-terminal amino acids of hFGF18. The RXXR motif is underlined, and the potential cleavage sites are shown as vertical arrows. **b,** Reactivity of 6 rabbit monoclonal antibodies to hFGF18. Purified human FGF181ΔC (hFGF181ΔC) from HEK293 cells was serially diluted and coated onto a plate. The coated hFGF181ΔC8 was detected with 1 μg/mL biotinylated 6 rabbit monoclonal antibodies followed by incubation with streptavidin-HRP. **c**, Comparison of the binding activities of clones 12S-SE5 and 12G7-9 and a commercially available rabbit polyclonal antibody (PA120525) to hFGF18. Purified hFGF181ΔC was coated and detected with 1 μg/mL of the indicated antibodies as described in **b. d**, Clone 19S-SE5 less efficiently binds to mouse FGF18 (mFGF18). Recombinant full-length mFGF18 was serially diluted and coated onto a plate. The coated proteins were detected as in **b**. Results are representative of two independent experiments.

Hence, we employed the binding capacity of hFGF18 to heparan sulfate to isolate hFGF18, which likely lacked the C-terminal amino acids. We utilized heparin columns to purify hFGF181ΔC using culture supernatants from HEK293 cells transiently expressing hFGF181ΔC. Then, we immunized rats and rabbits with purified hFGF181ΔC according to a standard procedure. We obtained 6 rabbit monoclonal antibodies and one rat monoclonal antibody (12G7-9). Since 6 rabbit mAbs showed similar binding activity to hFGF181ΔC (Fig. 1b), we used clone 19S-SE5 for subsequent experiments.

We coated serially diluted purified hFGF181ΔC and detected them using clones 19S-SE5 and 12G7-9 and a commercially available rabbit polyclonal anti-hFGF18 antibody (Cusabio CSB-PA120525). As shown in Fig. 1c, clone 19S-SE5 exhibited the highest efficiency in detecting coated hFGF181ΔC, albeit the signals of clones 12G7-9 and PA120525 were relatively low compared to clone 19S-SE5. Moreover, these three antibodies detected full-length mouse FGF18; the optical density (OD) of clones 12G7-9 and PA120525 reached a plateau at approximately 3 ng/mL of mFGF18. In contrast, the OD of 19S-SE5 still declined toward 0.1 ng/mL hFGF181ΔC, suggesting that the binding affinity of clone 19S-SE5 to mFGF18 was lower than that of hFGF181ΔC (Fig. 1d).

### 3.2. 19S-SE5 specifically recognizes the C-terminal portion of hFGF18

Assuming that the N-terminal signal peptides and C-terminal 5 to 7 amino acids may be deleted from FGF18 during maturation, the amino acid alignment of human and mouse FGF18 revealed that only one amino acid positioned at 183 was different: proline in hFGF18 versus alanine in mFGF18 (Fig. 2a). To define the epitope recognized by clone 19S-SE5, we used C-terminal truncated hFGF18 that lacked 8 amino acids (hFGF181ΔC8) and full-length mFGF18 produced in *E. coli* from commercial sources. The sensitivity of clone 19S-SE5 to mFGF18 was greatly reduced compared to hFGF181ΔC8 (Fig. 1c), suggesting that the primary epitope recognized by clone 19S-SE5 involves several amino acids, including proline^183^, at the C-terminus. We further deleted 25 amino acids, including proline^183^, to generate hFGF181ΔC25 in *E. coli* as a GST-fusion protein. As shown in Fig. 2b, the expression of hFGF181ΔC25 was verified by Coomassie blue staining. The deletion of the 25 C-terminal amino acids completely abolished the reactivity of clone 19S-SE5 (Fig. 2c), which suggests that the epitope recognized by clone 19S-SE5 is approximately 9 to 25 amino acids, including proline^183^ at the C-terminus of hFGF18. Deleting the 25 C-terminal amino acids reduced the activity of clone 12G7-9 toward hFGF181ΔC25, but clone 12G7-9 still weakly bound to hFGF181ΔC25 (Fig. 2d). These results suggest that the epitope recognized by clone 12G7-9 does not completely overlap with that of clone 19S-SE5.

**Fig. 2.**
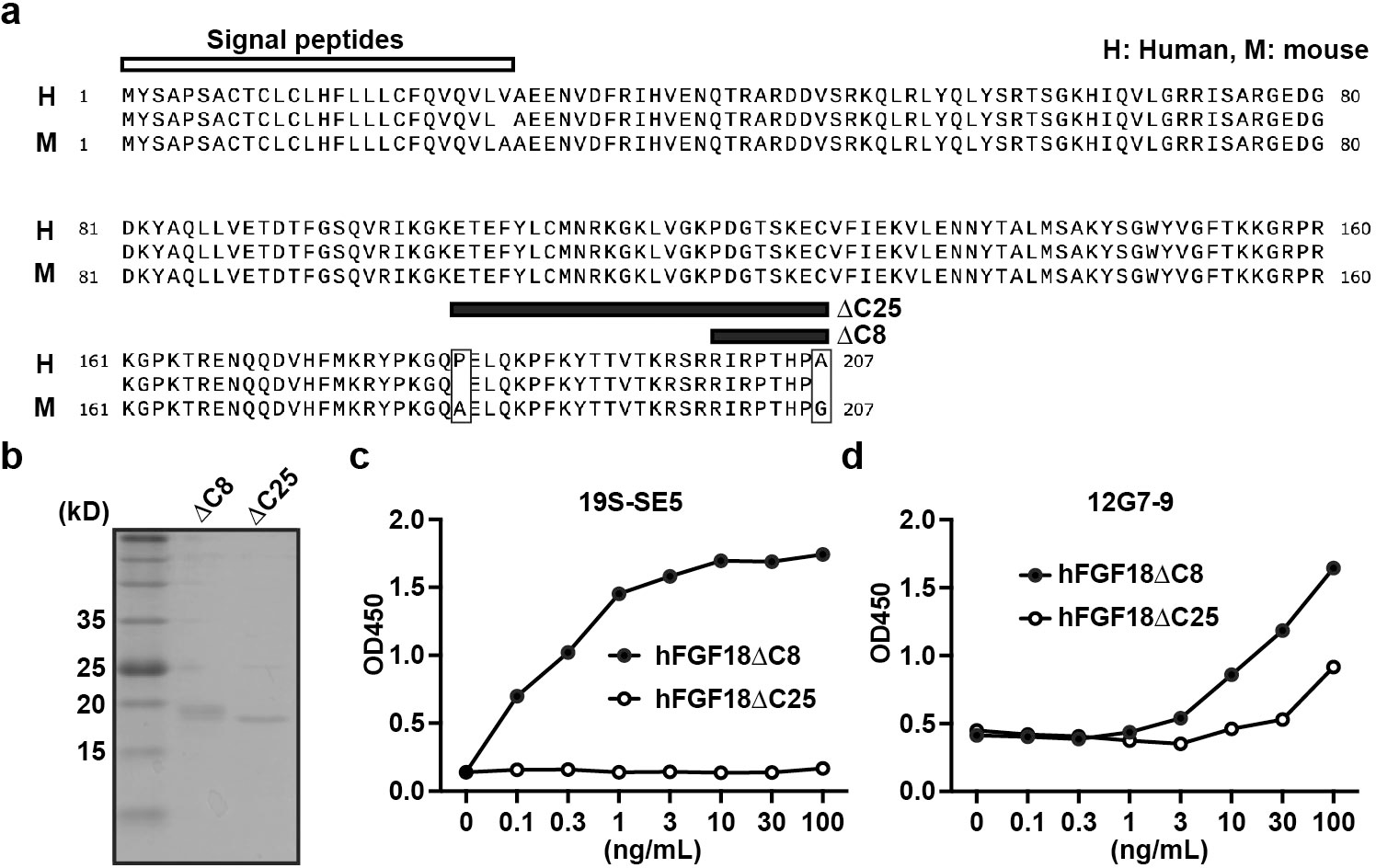
19S-SE5 specifically detects the C-terminal region of hFGF18. **a**, Alignment of human and murine FGF18 at protein levels. The first 26 amino acids indicate the signal peptides. 1ΔC8 and 1ΔC25 indicate the deleted amino acids from the C-terminus. The boxes indicate different amino acids between humans and mice. **b**, Coomassie blue staining of hFGF181ΔC8 and hFGF181ΔC25. GST-hFGF181ΔC25 was generated and purified in *E. coli* as described in the Materials and Methods section. Five micrograms of hFGF181ΔC8 and hFGF181ΔC25 were subjected to SDS‒PAGE and stained with Coomassie blue. **c, d**, Deletion of the last 25 amino acids completely and partially abolishes the binding activities of clones 19S-SE5 (**c**) and 12G7-9 (**d**), respectively. Purified hFGF181ΔC8 and hFGF181ΔC25 were serially diluted and coated onto the plate. The signals were detected as in Fig. 1b. Results are representative of two independent experiments.

### 3.3. Development of a high-sensitivity enzyme-linked immunosorbent assay (ELISA) for hFGF18

We developed a high-sensitivity ELISA using clone 12G7-9 as the capture antibody and clone 19S-SE5 as the detection antibody (Fig. 3a). Our assay detected hFGF18 at concentrations as low as 10 pg/mL.

**Fig. 3.**
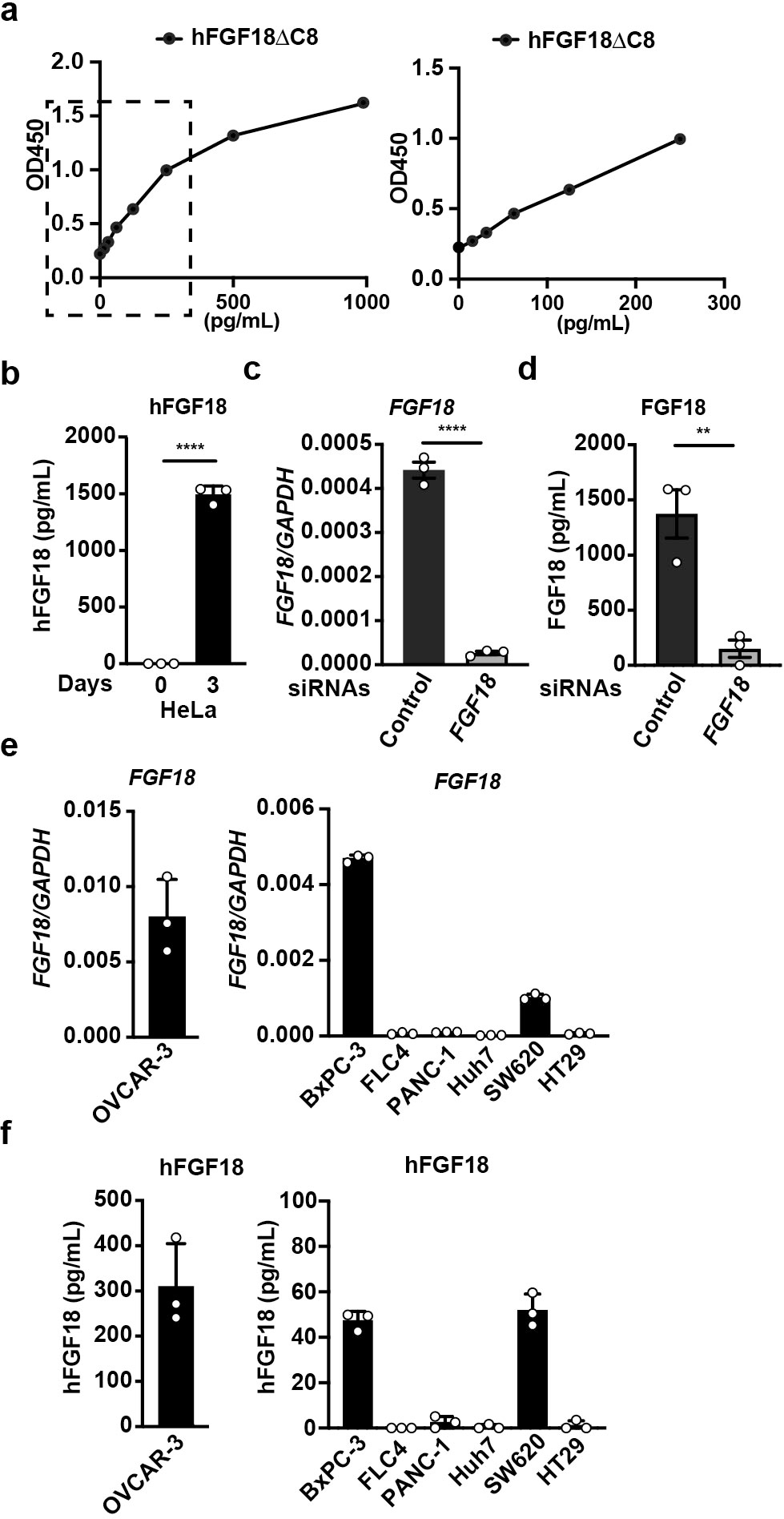
A sandwich ELISA to detect hFGF18. **a**, An in-house ELISA detecting human FGF18. The plate was precoated with clone 12G7-9, and then serially diluted hFGF181ΔC8 was added. The signals were detected with biotinylated clone 19S-SE5 and visualized by streptavidin polyHRP80, which enhanced the signals. The right panel is the enlarged image of the dotted box in the left panel. **b**, Expression of FGF18 is elevated in HeLa cells at protein levels. HeLa cells were cultured for 3 days, and the concentrations of FGF18 in the culture supernatants were determined by ELISA. Results are mean ± SD of triplicate samples. **c, d,** Knockdown of *FGF18* abolishes the expression of FGF18 in HeLa cells at both mRNA and protein levels. HeLa cells were transfected with control or *FGF18* siRNA and cultured for 3 days. The expression of *FGF18* was determined by qPCR (**c**). The concentrations of FGF18 in the culture supernatants were determined by an in-house ELISA (**d**). Results are mean ± SD of triplicate samples. **e**, Expression of *FGF18* mRNA in various tumor cell lines. The expression of *FGF18* was determined by qPCR. OVCAR-3, ovarian cancer; BxPC-3, pancreatic cancer; PANC-1, pancreatic cancer; Huh7, hepatocellular carcinoma; SW620, colorectal cancer; HT29, colorectal cancer. **f**, The concentrations of hFGF18 in the culture supernatants were determined by an in-house ELISA. The tumor cell lines were cultured for 3 days, and the concentrations of FGF18 were determined as in **b**. Results are mean ± SD of triplicate samples. Statistical significance was determined using a two-tailed unpaired Student’s *t-*test (**b, c,** and **d**). Results are representative of two independent experiments.

Previous studies reported that the expression of *FGF18* is elevated in the tumor tissues of several cancer patients [10,11,16]; hence, we measured the concentrations of FGF18 in culture supernatants from tumor cell lines. An in-house hFGF18 ELISA detected FGF18 in the culture supernatants from HeLa cells (Fig. 3b). Moreover, the knockdown of FGF18 expression by siRNA decreased the production of FGF18 in the culture supernatants and the mRNA expression of *FGF18* by qPCR (Fig. 3c, d), which indicated that the in-house ELISA system specifically detected hFGF18. To extend our analysis to other tumor cell lines, we measured the mRNA expression of seven cell lines, including OVCAR-3, BxPC-3, FLC4, Panc-1, Huh7, SW620, and HT29 (Fig. 3e). We detected FGF18 in the culture supernatants of OVCAR-3, BxPC-3, and SW620 cells (Fig. 3f), although the production of FGF18 was relatively low in BxPC-3 and SW620 cells.

### 3.4. Sensitivity comparison of the in-house ELISA to measure hFGF18 and mFGF18

The C-terminal portions of mFGF18 were almost identical to those of hFGF18 (Fig. 2a); therefore, it is plausible that several amino acids at the C-terminus should be cleaved during the maturation of mFGF18. To determine whether the deletion of these 8 amino acids could affect the reactivity of the in-house ELISA, we generated a C-terminal deletion mutant of mFGF18 lacking 8 amino acids (mFGF181ΔC8) in *E. coli*. We also generated full-length mFGF18 in *E. coli* as a control. The expression of these recombinant proteins was verified by Coomassie blue staining (Fig. 4a). We found that the in-house ELISA could detect mFGF18 irrespective of the presence or absence of the C-terminal 8 amino acids (Fig. 4a). Intriguingly, the reactivity toward mFGF181ΔC8 was higher than that observed for full-length mFGF18 (Fig. 4b), suggesting that the C-terminal 8 amino acids may mask the epitope that is recognized by the in-house ELISA. Of note, the reactivity of the in-house ELISA toward mFGF181ΔC8 was relatively low compared to hFGF181ΔC8; the lower detection limits of mFGF18 were approximately 500 pg/mL (Fig. 4c). Despite the relatively low sensitivity, we detected mFGF18 in sera from *Fgf18* Tg mice (Fig. 4d), which suggests that the in-house FGF18 ELISA would also be useful for measuring concentrations of mFGF18 in various murine experiments.

**Fig. 4.**
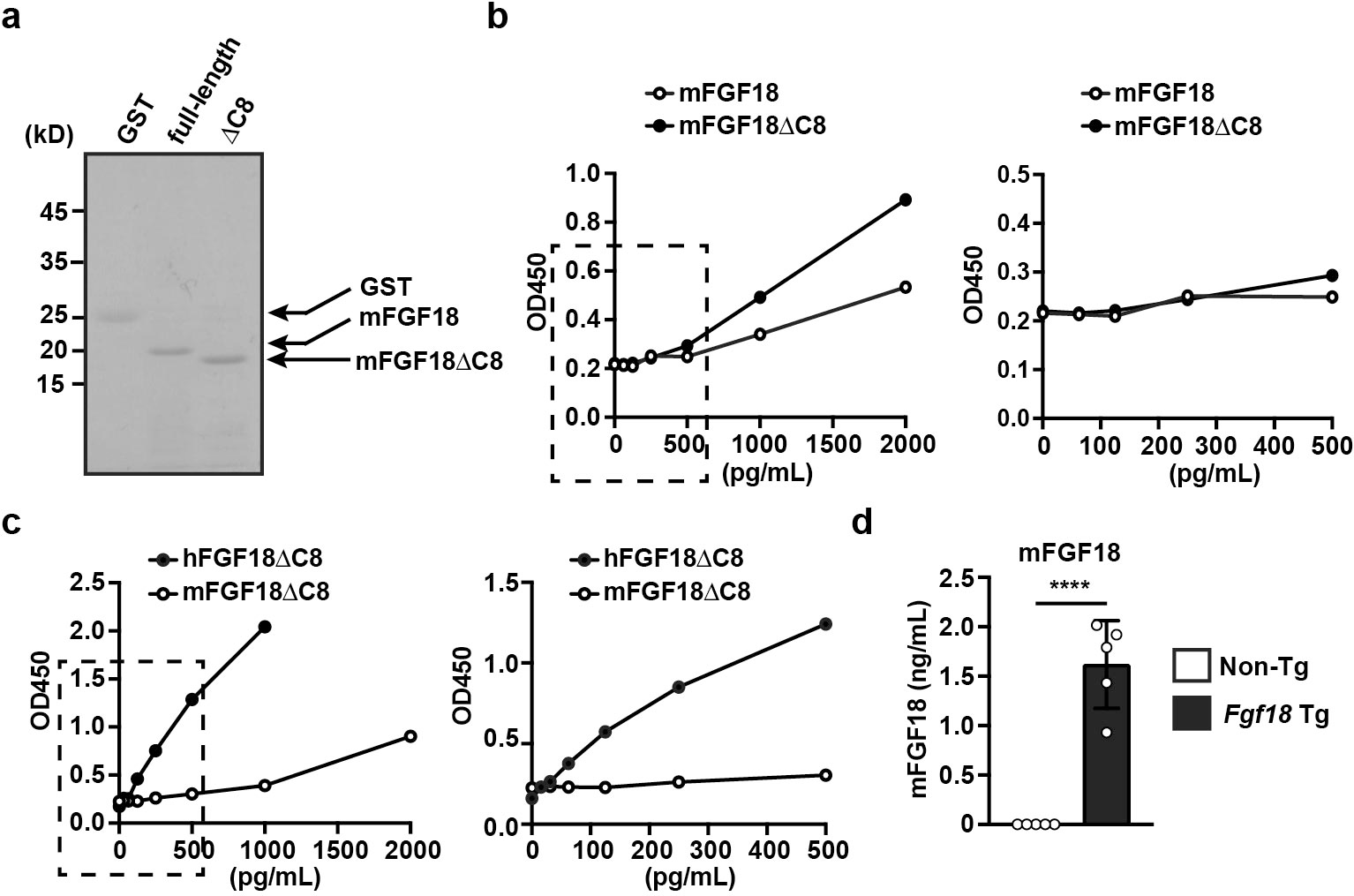
Comparison of the sensitivity of the in-house ELISA to detect hFGF18 and mFGF18. **a**, Coomassie blue staining of full-length mFGF18 and mFGF181ΔC8. GST-mFGF18 and GST-mFGF181ΔC8 were generated in *E. coli* and purified as in Fig. 2b. After digestion with PreScission protease, 5 μg of GST, mFGF18, and mFGF181ΔC8 were subjected to SDS‒PAGE and stained with Coomassie blue. **b**, Comparison of the sensitivity of the in-house ELISA to full-length mFGF18 and mFGF181ΔC8. The serially diluted proteins were coated, and the concentrations of the indicated proteins were determined by in-house ELISA. **c**, An in-house ELISA detected mFGF181ΔC8, albeit the sensitivity is relatively low compared to hFGF181ΔC8. The serially diluted hFGF181ΔC8 and mFGF181ΔC8 were coated, and the concentrations of hFGF181ΔC8 and mFGF181′8 were determined by the in-house ELISA. The right panel is the enlarged image of the dotted box in the left panel. Results are representative of two independent experiments. **d**, mFGF18 is elevated in sera from *Fgf18* Tg mice. Sera from 6- to 8-week-old Non-Tg and *Fgf18* Tg mice and concentrations of mFGF18 were determined by in-house ELISA. Results are the mean ± SE (*n* = 5 mice). Statistical significance was determined using a two-tailed unpaired Student’s *t-*test. Results were pooled from two independent experiments.

In this study, we developed a high-sensitivity ELISA for hFGF18 and detected hFGF18 in culture supernatants from several cancer cell lines. However, it is unclear whether the in-house ELISA could detect hFGF18 in sera from various cancer patients. Our preliminary study revealed that we could detect hFGF18 in sera from small numbers of cancer patients. To demonstrate the efficacy of FGF18 as a biomarker, additional investigations will be necessary, such as the enrollment of a substantial number of cancer patients and measurement of hFGF18 levels in their sera throughout cancer progression or before and after treatment. Although these experiments may be time-consuming, they have the potential to offer valuable insights into the clinical significance of FGF18.

## Acknowledgments

We thank E. Miyoshi and T. Natsume for providing the FLC4 cell line and pcDNA3-FLAG, respectively. This work was supported in part by Grants-in-Aid for Scientific Research (B) 20H03475 (to HN) and 23H02707 (to HN), Scientific Research (C) 17K08994 (to YT) and 20K11589 (to YT) from the Japan Society for the Promotion of Science (JSPS), the Japan Agency for Medical Research and Development through AMED-CREST (grant no. 22gm1210002; to MT and HN), Princess Takamatsu Cancer Research Fund (to HN), and Toho University Grant for Research Initiative Program (TU-GRIP) (to HN).

## Contributions

Y.T. and H.N. designed the study and interpreted the results. M.T., K.C.E., C.J.O., and T.M. generated antibodies against hFGF18. Y.T., S.K-S., T.K., and K.M. performed and analyzed most of the experiments. H.A., H.Y., and K.O. supervised the experiments. Y.T. and H.N. wrote the manuscript with constructive input from all authors.

## Data availability

The authors declare that the data supporting the findings of this study are available within the paper and its supplementary information files. Data not included are available from the corresponding authors upon reasonable request.

## Abbreviations

FGF: Fibroblast growth factor
ELISA: enzyme-linked immunosorbent assay.

